# Normative models of individualized functional brain networks reveal language network expansion in autism

**DOI:** 10.1101/2025.10.23.684286

**Authors:** Ruoqi Yang, Xinyu Wu, Shuo Lv, Jinlong Li, Zhiming Wang, Wenjing Zhu, Tan Gao, Guoyuan Yang

**Affiliations:** School of Interdisciplinary Science, Beijing Institute of Technology, Beijing, China; School of Medical Technology, Beijing Institute of Technology, Beijing, China

**Author notes:** Corresponding authors: Guoyuan Yang.

## Abstract

Autism spectrum disorder is a highly heterogeneous neurodevelopmental disorder, hindering mechanistic insights and the identification of biomarkers for clinical diagnosis. Recently, precision functional mapping has been developed to identify abnormalities in brain network topologies associated with various psychiatric disorders, yet its application in autism remains limited. Here, we utilized precision functional mapping and a large, multisite neuroimaging dataset (*N* = 1,182) to construct individualized functional networks in individuals with autism. We developed normative models using network surface area from healthy controls (n = 628) to characterize typical brain network organization across age, allowing for the quantification of individual-specific deviations in individuals with autism (n = 554). We found widespread and heterogeneous deviations from the normative model, with the language network emerging as the most significantly altered region, thereby emerging as an epicenter of functional disruption in autism. Individuals with autism were clustered into three subtypes involving distinct functional network topologies, associated with behavioral profiles marked by impairments in perception, language processing, or socio-emotional functioning. We further linked these atypical brain features to cortical gene expression patterns, revealing enriched pathways related to neurodevelopment, language, and signaling processes. Together, these findings reveal autism-specific deviations in individualized functional brain networks, offering potential clinical relevance for understanding and stratifying autism.

## Introduction

Autism spectrum disorder (ASD) is a major neurodevelopmental disorder, characterized by persistent deficits in social communication and interaction, alongside restricted and repetitive patterns of behavior^1,2^. There is considerable individual variability in both its clinical manifestations and the neural functional basis^3^. Decades of functional magnetic resonance imaging (fMRI) studies have tested comparisons between groups of ASD and healthy subjects^4^. Typically, previous studies focus on ASD at the level of functional connectivity^5–7^ and structural measures^8–10^ to elucidate the underlying neurophysiological substrates of the disorder. However, these findings exhibit significant inconsistencies, resulting in a lack of clear mechanistic insights regarding the neurophysiological mechanisms of ASD and making them difficult to generalize to individualized clinical reality. Recently, precision functional mapping has been applied to systematically characterize brain function and organization at the individual level, revealing that individual brain network topology exhibits substantial variability in size, shape, and spatial location compared to group averages^11–13^. Although there is currently one study on whole-brain functional parcellation in individuals with ASD, reporting reduced voxel-level stability and blurred network boundaries across multiple systems^14^, the small sample size makes it difficult to accurately define the normal range, and the lack of research on disease subtypes prevents the establishment of a universally applicable reference standard. This situation compels us to highlight the need for further investigation through large-scale, individual-level precision mapping studies of ASD.

The high heterogeneity of ASD generally leads to poor replication of neuroimaging findings across small datasets, posing a significant barrier to their translation into clinical diagnostic practice^15–19^. Recent pioneering work in systems neuroscience has begun to expand sample sizes across a broad age range^20,21^, along with techniques to harmonize multisite data^22^. Normative modeling is a statistical approach designed to characterize the range of normal variation in specific features, such as brain structure or function^20,23–28^. Unlike traditional case–control analyses, normative models aim to characterize the distribution of brain-related variables within typically developing populations and use this as a reference to quantify individual-level deviations, that has been increasingly applied to ASD research^8,29^. Normative modeling can incorporate demographic factors (e.g., age, sex) to account for individual variability in resting-state brain function, thereby offering a powerful framework to explore the clinical relevance of functional heterogeneity, as well as its potential associations with ASD subtypes^22,30^. Previous study has revealed that individuals with ASD exhibit highly individualized and widely distributed patterns of cortical thickness deviations and the spatial distribution of these deviations varying markedly across individuals^8^. Similarly, ASD individuals also demonstrate pronounced nonlinear and highly individualized deviations in the developmental trajectories of resting-state functional connectivity, particularly involving the default mode, somatomotor, and ventral attention networks, with these deviations closely associated with core ASD symptoms^29,31^. Although these findings provide important clues for understanding the neural mechanisms of ASD, they have largely overlooked the fact that functional brain network topology which can change significantly across the lifespan^21^, and showed behaviorally meaningful differences in brain network topography and connectivity in psychiatric disorders^13^. The construction of normative model based on large, age-diverse populations for individualized functional network topology, and quantitatively measured deviations from typical developmental trajectories of ASD, can provide ASD-specific insights into network topological variations and inform precision ASD subtypes.

ASD is a highly heritable neurodevelopment disorder, with twin and family studies estimating heritability rates reach to 80–90%^32,33^. Over the past decade, researchers have identified dozens of genes linked to increased susceptibility to ASD, with these genetic factors collectively accounting for 10–20% of ASD cases^34^. Many of the identified risk genes are involved in neurodevelopmental processes, such as synaptic function, neuronal migration, and chromatin remodeling^35,36^, suggesting that alterations in brain development is a key mechanism underlying ASD pathophysiology. However, existing studies have yet to investigate the underlying genetic mechanisms that may contribute to the aberrant topological organization of functional brain networks observed in individuals with ASD^14^. Previous studies have identified distinct heritability patterns underlying the characteristics of the brain’s functional topology^37,38^. Understanding how such systems-level brain aberrations link to specific genetic architectures could provide new insights into the biological basis of ASD heterogeneity^39^. Therefore, examining links between ASD-related topological alterations in brain functional networks and gene expression enables identification of spatially patterned molecular features that may underlie observed disease heterogeneity.

Here, we applied a normative modeling approach by incorporating individualized precision functional maps to better understand the neural heterogeneity of ASD using a multisite resting-state fMRI dataset of 1,182 male participants from the Autism Brain Imaging Data Exchange (ABIDE) I and II datasets. This shifts the focus away from group-level comparisons toward characterizing the degree of alteration in each individual with reference of the healthy controls (HCs) ^8^, allowing us to identify meaningful individual-level deviations from the normative range and account for the relationship between brain functional regions and behavioral performance in ASD^40^. To achieve this, we first delineated the topology of individual functional brain networks and quantified the surface area percentage of each functional network (Fig. 1a). Then, we constructed normative models using data from HCs to estimate the expected range of surface area distribution across brain networks. Our main goals were to (1) identify cortical regions where ASD individuals show significant deviations from normative models, defining core neurofunctional epicenters of abnormality (Fig. 1b); and thereby (2) to stratify ASD into subtypes based on individualized patterns of functional network topology deviations and link these subtypes to distinct behavioral profiles (Fig. 1c); and finally (3) to explore the potential genetic underpinnings of these atypical functional patterns (Fig. 1d).

**Fig. 1.**
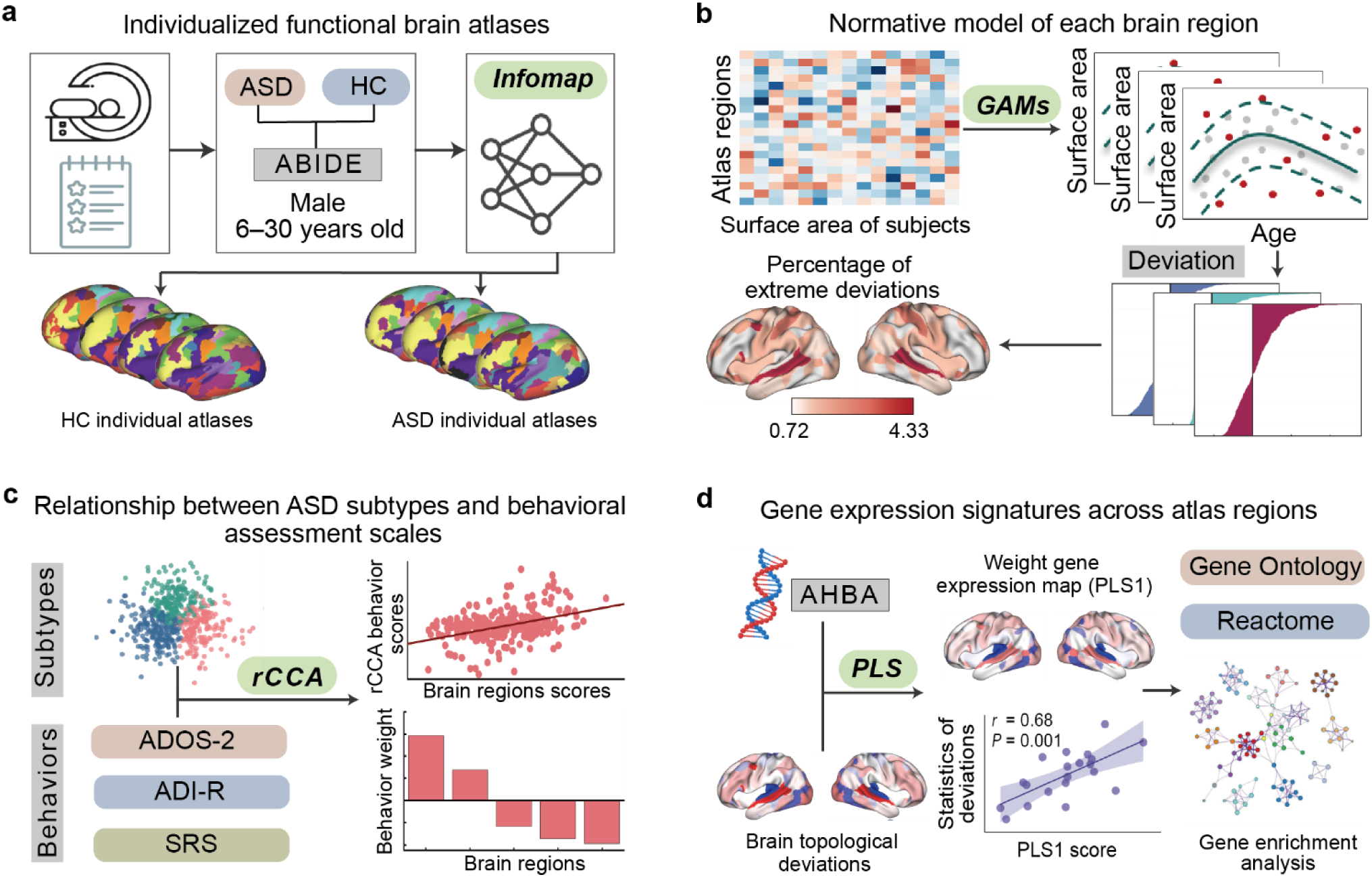
Overview of the analytic workflow. **a** Individualized functional brain atlases were generated for autism spectrum disorder (ASD) and healthy control (HC) participants using resting-state fMRI data from the ABIDE I and II databases^109,110^ and the Infomap community detection algorithm^12,13^. **b** Generalized additive models (GAMs) were applied to HC participants data to establish age-related normative models for the surface area of each brain region. Deviation scores for ASD individuals were calculated by comparing their regional surface areas to the normative trajectories. The overlap brain map shows where individuals with extreme deviations are most concentrated across brain networks. **c** Regularized canonical correlation analysis (rCCA)^124,125^ was used to examine the relationships between ASD subtypes identified through clustering and behavioral measures, including the Autism Diagnostic Observation Schedule - Second Edition (ADOS-2)^114^, Social Responsiveness Scale (SRS)^115^, and Autism Diagnostic Interview - Revised (ADI-R)^116^ behavioral scales. **d** Partial least square regression (PLS) was performed to relate brain region deviations to gene expression data from the Allen Human Brain Atlas (AHBA)^128^. The topographical PLS1 gene expression map was correlated with surface area differences between ASD and HC participants. Gene enrichment analyses were conducted using Gene Ontology^69^ and Reactome databases^70^ to interpret the biological relevance of the identified genes.

## Results

### Individualized functional mapping reveals topographic heterogeneity in autism

We first generated individualized precision functional maps for all participants with ASD and HCs from the ABIDE I and II database with the age range from 6 to 30 years (Table 1), and examined inter-individual variability in cortical functional networks. Functional networks were identified using the Infomap community detection algorithm, which detects modules based on patterns of spontaneous functional connectivity across the cortical parcels at the individual level^12,13^. To ensure methodological consistency, a common group-level prior template was used for both ASD and HC groups^13^. We applied ComBat harmonization to the surface area percentages for each network to reduce non-biological variability introduced by site-related factors, while preserving meaningful biological differences^41,42^. After harmonization^43^, site effects which assessed using analysis of variance (ANOVA) showed not significant for each functional brain network (Supplementary Fig. 1). Then, the group-level atlases for ASD and HC groups were derived by assigning each vertex to the network whose label had the highest mode across individual assignments. In the group level, individuals with ASD and HCs exhibited largely similar spatial arrangements of each cortical functional network (Fig. 2a), suggesting that group-averaged analyses may obscure individual-level heterogeneity within the ASD population. Compared to group-averaged atlases, individualized functional brain atlases revealed pronounced heterogeneity in the spatial arrangement and boundaries of several cortical networks, particularly in the parietal and dorsolateral subdivisions of the default mode network, visual, somatomotor, and especially language networks (Fig. 2b), indicating that these networks exhibited the most substantial individual-level variability. Importantly, this expansion does not necessarily reflect enhanced language function; rather, it may indicate atypical developmental trajectories of the language network and special communication patterns in individuals with ASD^44,45^, leading to altered cortical parcellation boundaries or compensatory functional reorganization.

**Fig. 2.**
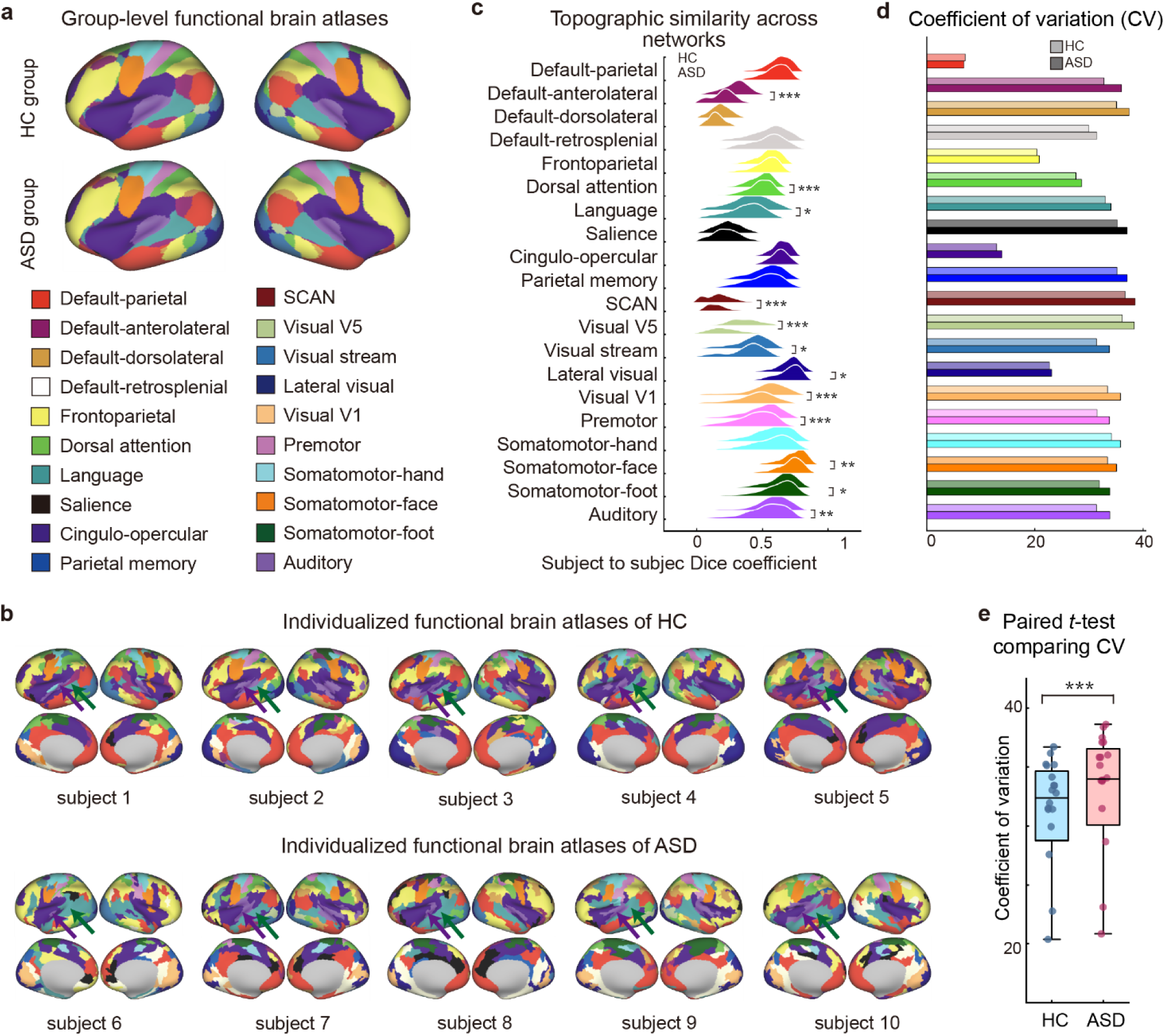
Group-level and individualized functional brain atlases in ASD and HC participants. **a** Group-level functional brain atlases for HC individuals (top panel) and individuals with ASD (bottom panel), displayed on inflated views of the cortical surface using Connectome Workbench (http://www.humanconnectome.org/software/connectome-workbench). **b** Example of individualized functional brain networks in the cortex for HCs and individuals with ASD. The language (green arrows) and auditory (purple arrows) networks are highlighted to illustrate the variations of network topology in cortical territory. **c** The ridge plot displays distributions of inter-subject Dice coefficients across each network, with top ridge represents HCs and bottom ridge represents ASD individuals (KS test, \**P* < 0.05, \*\**P* < 0.01, \*\*\**P* < 0.001, FDR corrected). **d** Variation of inter-subject similarity in Dice coefficients across 20 networks, measured with coefficient of variation (CV), which corrects for differences in the total average size of each network. **e** Boxplot of CV across 20 networks for HCs and individuals with ASD after regressing out group-average network size (paired *t*-test; *t* = 8.868, *P* < 0.001). SCAN, somato-cognitive action network.

**Table 1.**
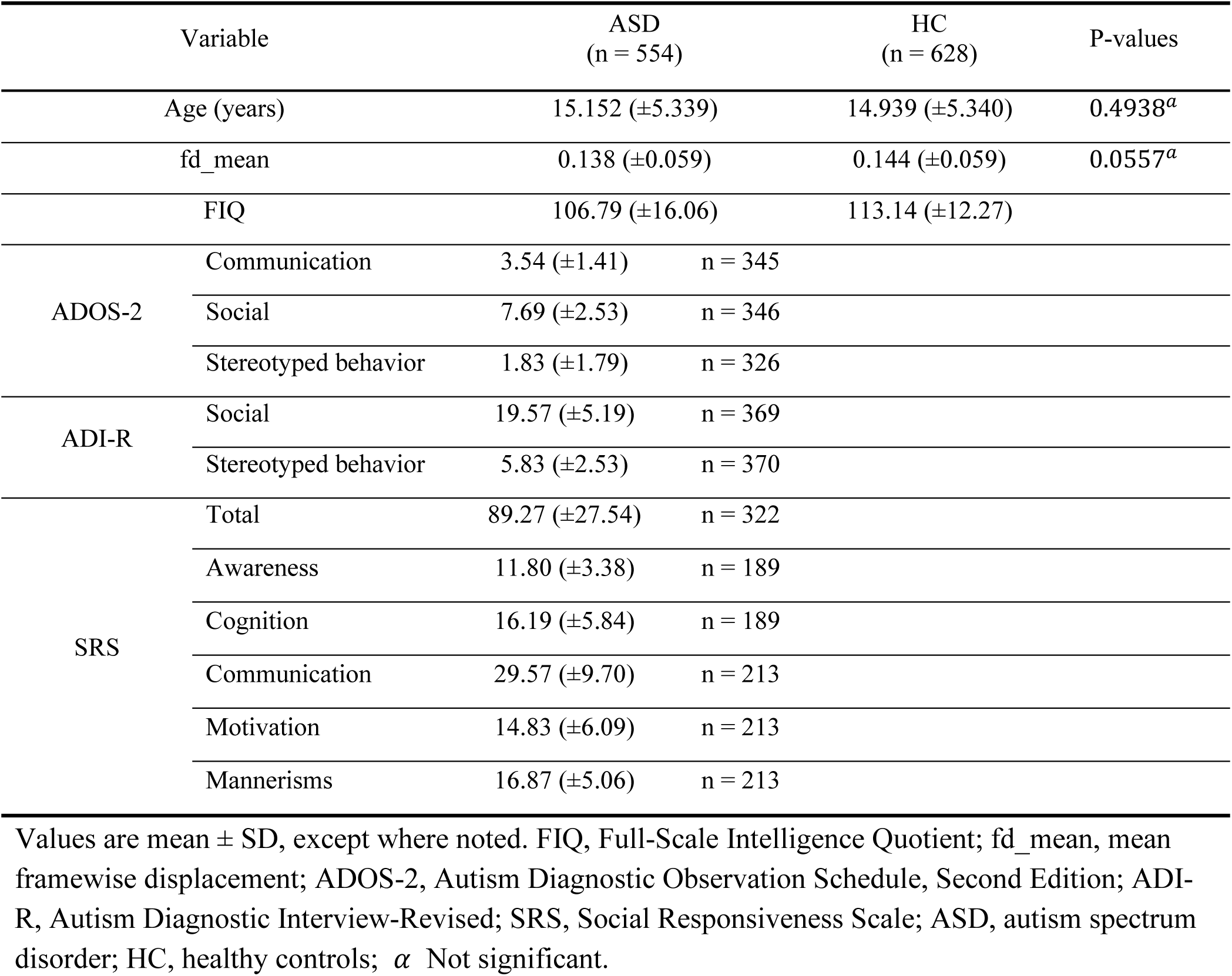
Demographic and diagnostic characteristics of study participants.

To quantitatively assess the consistency of functional network topography across individuals, we further computed the subject-to-subject Dice coefficient across 20 canonical brain networks in both ASD and HC groups (Fig. 2c). We observed that primary sensory and motor networks, including lateral visual, somatomotor-face, and auditory networks, exhibited high inter-individual similarity in both groups, reflecting robust and conserved spatial organization. In contrast, association networks— particularly those involved in higher-order cognition such as the default mode, frontoparietal, and language networks—showed markedly reduced Dice similarity in both ASD group and health controls^37^. Compared to HCs, participants with ASD exhibited greater inter-individual variability, suggesting disrupted topographic consistency in these networks among individuals with ASD. We further performed Kolmogorov–Smirnov (KS) tests across individual Dice similarity distributions for each network^46^, and the results revealed that 12 out of 20 networks showed significant group differences (all *P*-values < 0.001, KS test with FDR corrected). These findings shows that both primary sensory cortices and transmodal association systems may be differentially impacted in ASD, exhibiting altered topographic consistency relative to HCs. In addition, the pronounced heterogeneity in functional network topology among individuals with ASD is further demonstrated by the coefficient of variation (CV) across 20 networks after regressing out group-average surface area size for each network (Fig. 2d). Overall, ASD group shows significantly greater variation of inter-individual similarity than HC groups, reflecting increased diversity in functional network topology (Fig. 2e; *t* = 8.868, *P* < 0.001). This heightened variability may indicate systematic abnormalities in the functional brain topology associated with ASD, potentially resulting in impairments in language and social communication abilities in individuals with ASD^47,48^.

### Individualized deviations from normative developmental trajectories in ASD

The results above indicate that topographic features of ASD groups exhibited more inter-individual variability compared to HCs. To quantify this variability for each network, we further applied normative modeling frameworks to calculate atypical deviations in individuals with ASD. Ten-fold cross-validated generalized additive models (GAMs) were trained on HCs to model the age-related trajectory of cortical surface area for each functional network^20^. Individual-level deviations were then derived by projecting each individual’s surface area percentages across all functional networks, aligned with their age, onto the normative developmental trajectories derived from the HCs. We first applied principal component analysis (PCA) to summarize the multivariate structure of these deviations across all networks (Fig. 3a). The first principal component differentiated ASD from HC groups, indicating the presence of systematic divergence from typical development in participants with ASD. Notably, ASD group exhibited greater dispersion along principal component 1, reflecting marked within-group heterogeneity. Deviations above the normative trajectories, indicating that an individual’s surface area percentages exceeded the age-matched typical developmental average, were defined as positive deviations, while those below were defined as negative deviations^8^. To statistically assess group differences in these deviations, we performed two-sample *t*-tests for each network. The analysis revealed that four networks showed statistically significant deviations in participants with ASD (*P* < 0.05, FDR-corrected), including the language, auditory, visual V5, and somatomotor-foot networks (Fig. 3b). These significant deviations were predominantly localized to the temporal, insula and posterior occipital cortices (Fig.3c). Figure 3d shown individuals with ASD exhibited marked deviations from normative surface area trajectories across these four significantly affected networks. Positive deviations were most prominent in the language network. In contrast, consistent negative deviations were observed in the auditory, visual V5, and somatomotor-foot networks. For other brain networks in individuals with ASD that deviate from the normative model, please refer to the Supplementary Figure 2. The normative deviation distributions across distinct brain regions in individuals with ASD demonstrate region-specific biases (Fig. 3e). For the distribution of deviation scores of other networks, please refer to the Supplementary Figure 3. Notably, the language network exhibited the highest proportion of extreme positive deviations and it is accompanied by the extraction of the auditory network, highlighting the pivotal role of language-related regions in the neurodevelopmental alterations associated with ASD. Meanwhile, negative deviations in the visual V5 and somatomotor-foot networks may impair the integration of motion signals and provide a neural basis for the difficulties participants with ASD experience in perceiving and interpreting dynamic visual information^49,50^.

**Fig. 3.**
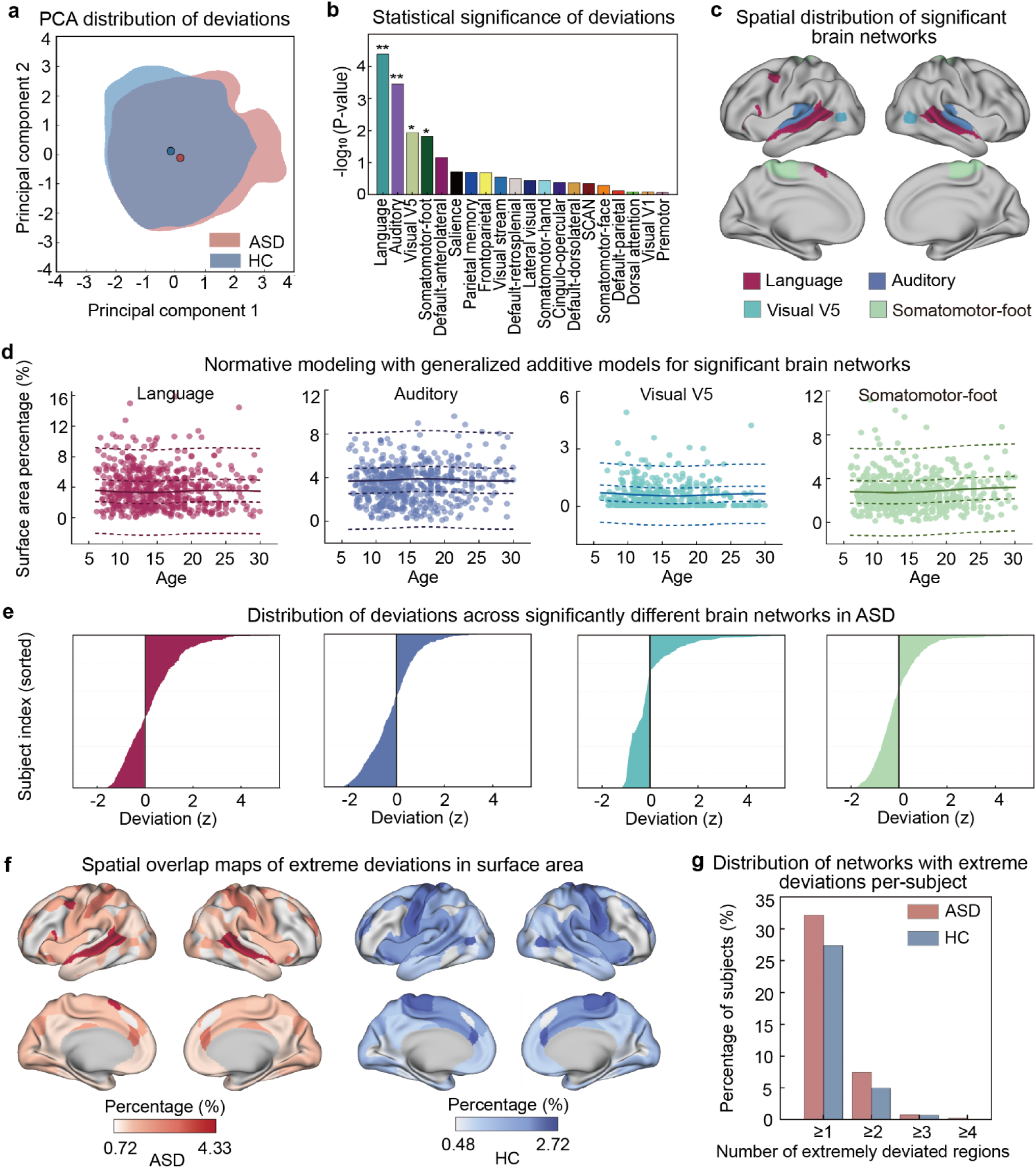
Normative models established in HC participants and deviations from normative models in individuals with ASD. **a** Overlap map depicts the distribution of deviations in the space of the first and second principal components (PC1 and PC2) for the ASD (red) and HC (blue) groups. Shaded areas show group distribution ranges, and dots represent their central points. **b** Bar plot depicting the -log_10_ (*p*-value) of brain networks with significant surface area deviations in individuals with ASD relative to normative models. Asterisks signify statistical significance with FDR corrected q < 0.05 (\**P* < 0.05, \*\**P* < 0.01). **c** Brain networks with significant surface area deviations in ASD individuals compared to HCs across cortex, with negative deviations (blue, purple, green) and positive deviations (magenta). **d** Normative models from HCs in each brain network (language, auditory, visual V5 and somatomotor-foot) established by GAM. The median (50^th^) centile is represented by a solid line, while the 5^th^, 25^th^, 75^th^ and 95^th^ centiles are indicated by dashed lines. Each scatter point represents the surface area percentage value of an individual with ASD in the corresponding brain network. **e** Bar chart shows the distribution of deviation scores (in z-scores) for individuals with ASD. **f** The spatial overlap maps illustrate the percentage of patients with extreme deviations from the normative range for each brain network at a threshold of z > |2.6| (left panel for ASD group; right panel for HC group). **g** Bar plots show the number of networks per-subject with extreme deviation regions in individuals with ASD (red) and HCs (blue).

We further verified the cortical spatial distribution characteristics of network deviations in participants with ASD and the differential patterns of HCs by calculating the spatial overlap maps of extreme deviations. These deviations were defined based on a threshold of |z| ≥ 2.6, corresponding to a 99% confidence interval^20,51,52^. The spatial overlap maps showed that individuals with ASD exhibited a higher overall frequency of extreme deviations, which were primarily concentrated in temporal and parietal cortices, with most observed in the language network (Fig. 3f). In contrast, extreme deviations in HCs were distributed sparsely and uniformly across brain networks with somatomotor and cingulo-opercular networks showing a higher overlap. In addition, the number of extreme deviation networks per participant revealed that, compared to HCs, a larger proportion of individuals with ASD had networks showing extreme deviations (Fig. 3g). Together, these findings suggest that network-level alterations in ASD affect both higher-order associative systems and lower-level sensorimotor systems, particularly in the language network, which may play a key role in the neurodevelopmental trajectory of ASD.

### Network topology deviation-based ASD subtypes associated with different behavioral domains

Previous studies have widely employed neurosubtyping approaches to characterize ASD, yet findings remain inconsistent across investigations, largely attributable to inadequate accounting for underlying neurobiological heterogeneity^18^. Here, based on deviation features from normative models, we clustered individuals with ASD into three functional subtypes (Fig. 4a). These three subtypes exhibited heterogeneous deviation patterns across cortex, with each subtype represents a unique pattern of functional disruption within ASD populations (Fig. 4b). Specifically, subtype 1 was characterized by enlargement in the lateral visual and default-parietal networks, and reduction in the auditory, visual stream, visual V1 and premotor networks (Supplementary Fig. 4a). Subtype 2 primarily involved abnormalities in the language network (Supplementary Fig. 4b), while subtype 3 showed moderate but widespread deviations across multiple brain networks, particularly in the visual network (Supplementary Fig. 4c).

**Fig. 4.**
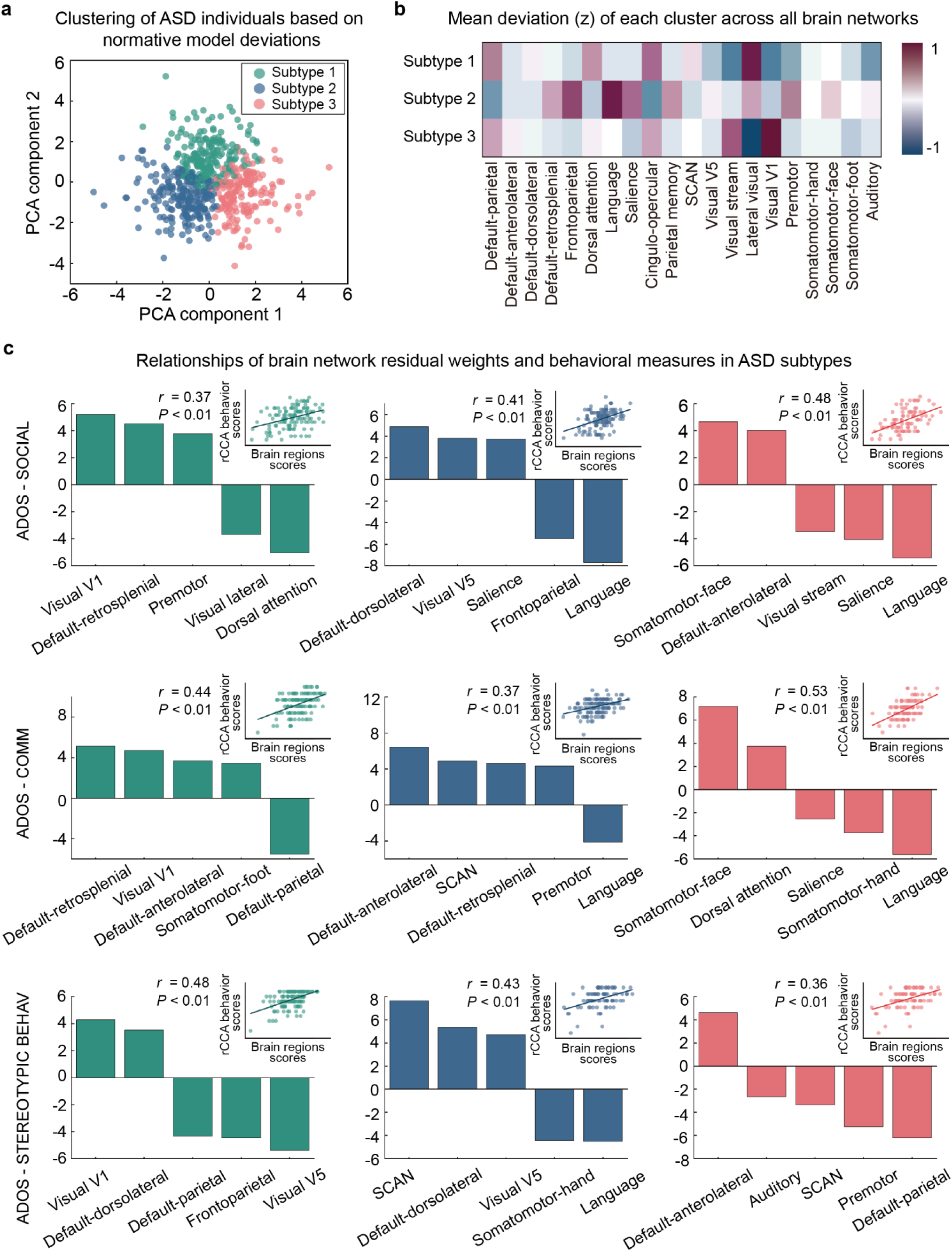
The relationship between brain network topological deviations and behavioral measures in ASD subtypes. **a** The scatter plot illustrates the clustering results of individuals with ASD using k-means clustering algorithm based on deviations from normative models. Different colors denote three identified ASD subtypes: subtype 1 (green), subtype 2 (blue), and subtype 3 (red). **b** The heatmap displays the mean deviation (z-scores) of each ASD subtype across various brain networks. **c** The relationships between the top 5 weights of brain networks and ASD-related behavioral measures across three subtypes. The bar charts display the weight values corresponding to brain networks and behaviors, including the ADOS - SOCIAL (Autism Diagnostic Observation Schedule - Social domain), ADOS - COMM (Communication domain), and ADOS - STEREOTYPIC BEHAV (Stereotypic Behavior domain). The scatter plots represent the correlations between brain regions topological deviations and behavioral measures derived from rCCA.

We then examined the associations between functional network deviation patterns and clinical behaviors across subtypes, using measures from various subscales of the Autism Diagnostic Observation Schedule, second edition (ADOS-2), through regularized canonical correlation analysis (rCCA) (Fig. 4c). We found that in subtype 1, social and communication deficits were associated with positive CCA weights in the primary visual cortex (V1), default-retrosplenial, and premotor networks, and negative weights in the dorsal attention, lateral visual and default-parietal networks (social: *r* = 0.37, communication: *r* = 0.44, all *P*-values < 0.01). Stereotyped behaviors were correlated with positive weights in the visual V1 and default-dorsolateral networks, and negative weights in the frontoparietal, visual V5, and default-parietal networks (*r* = 0.48, *P* < 0.01). Thus, individuals in this subtype may experience difficulties in processing socially relevant perceptual cue, such as facial expressions and nonverbal signals like gestures, and may exhibit more pronounced stereotyped behaviors due to reduced cognitive control^53,54^. In subtype 2, difficulties in social and communication functioning were associated positive canonical weights in higher-order cognitive and executive networks, including the default-anterolateral, default-dorsolateral, somato-cognitive action (SCAN), and dorsal attention networks, and negative weights in the language and frontoparietal networks (social: *r* = 0.41, communication: *r* = 0.37, all *P*-values < 0.01). Stereotyped behaviors were positively linked to the SCAN, default-dorsolateral and visual V5 networks, and negatively associated with the language and somatomotor-hand networks (*r* = 0.43, *P* < 0.01). The social and language impairments observed in this subtype may stem from deficits in information integration, contextual understanding, and behavioral planning^54–56^. In subtype 3, both social and communication difficulties were associated with positive canonical weights in the somatomotor-face and attention-related networks, with the default-anterolateral network contributing to social difficulties and the dorsal attention network contributing to communication difficulties. In contrast, negative canonical weights were observed in the salience and language networks (social: *r* = 0.48, communication: *r* = 0.53, all *P*-values < 0.01), suggesting increased engagement of face-processing and attentional systems, potentially reflecting compensatory mechanisms, alongside reduced integration of the language, salience, and sensorimotor networks. Additionally, stereotyped behaviors were associated with negative canonical weights in the auditory, premotor, SCAN, and default-parietal networks (*r* = 0.36, *P* < 0.01), reflecting reduced functional integration across the somatomotor and default mode networks. These findings suggest that subtype 3 may represent an ASD phenotype characterized by core socio-emotional dysfunction^57^. Behavioral impairments in this group are likely rooted in difficulties with emotion recognition, affective communication, and sensory input processing^58^.

Furthermore, we conducted replicated analyses of the behavioral association patterns using additional clinical behavioral scales. We found that these three functional subtypes exhibited distinct brain–behavior associations, consistently observed across both the Autism Diagnostic Interview-Revised (ADI-R) (Supplementary Fig. 5) and the Social Responsiveness Scale (SRS) scales (Supplementary Fig. 6). Each subtype showed unique regional patterns of association with core symptom domains, including social communication, repetitive behaviors, social awareness, cognition, and behavioral regulation, further reinforcing the neural specificity identified in the above analyses. Specifically, subtype 1 was primarily associated with perceptual and sensorimotor regions, subtype 2 with executive and language-related networks, and subtype 3 with systems involved in affective processing and sensory integration.

To evaluate if individualized deviation patterns can predict the subtypes of participants with ASD, we trained a variety of supervised machine models with deviation of surface area in each functional network across three ASD subtypes (Supplementary Fig. 7a). These include random forest (RF), support vector machine (SVM), and logistic regression (LR). Specifically, individualized network deviation values for all networks were given as input for model training (20 total features). We employed a standard 5-fold cross-validation scheme to optimize model parameters (see the *Methods* for more details). Supplementary Figure 7b shown that the area under the receiver operating characteristics curve (AUC) for LR was highest (AUC = 0.96), followed by SVM (AUC = 0.95), and RF (AUC = 0.91). Similarly, the confusion matrix indicates that the multiclass classifier exhibits higher certainty in identifying each ASD subtypes (Supplementary Fig. 7c). Together, these results indicate that the deviations in whole brain network topology in participants with ASD can serve as effective features for identifying its corresponding subtypes, suggesting that the network topology deviation patterns of the normative models have the potential to be used in the subtype classification of mental illness.

### Genetic signatures underlying cortical topology deviations in ASD

To further investigate the associations between ASD-specific network topological characteristics and genetic signatures, we applied partial least squares (PLS) regression between the statistics map derived from surface area deviations in participants with ASD and Allen Human Brain Atlas (AHBA, http://human.brain-map.org) cortical gene expression patterns^59^. The first component (PLS1), which captures the most significant association between gene expression data and ASD predictive statistic of deviations (Fig. 5a, top panel), explained 38.8% of the total variance in the response variables (Supplementary Fig. 8a), with significance confirmed by a nonparametric permutation test (Supplementary Fig. 8b; *P* < 0.001). We observed that the PLS1 weighted map was spatially correlated with the statistic of deviations (Fig. 5a, bottom panel; *r* = 0.63, *P* = 0.003). This positive correlation indicates that genes with positive (e.g., language network) or negative (e.g., auditory network) PLS1 weights tend to be overexpressed in regions showing corresponding positive or negative ASD effects, respectively.

**Fig. 5.**
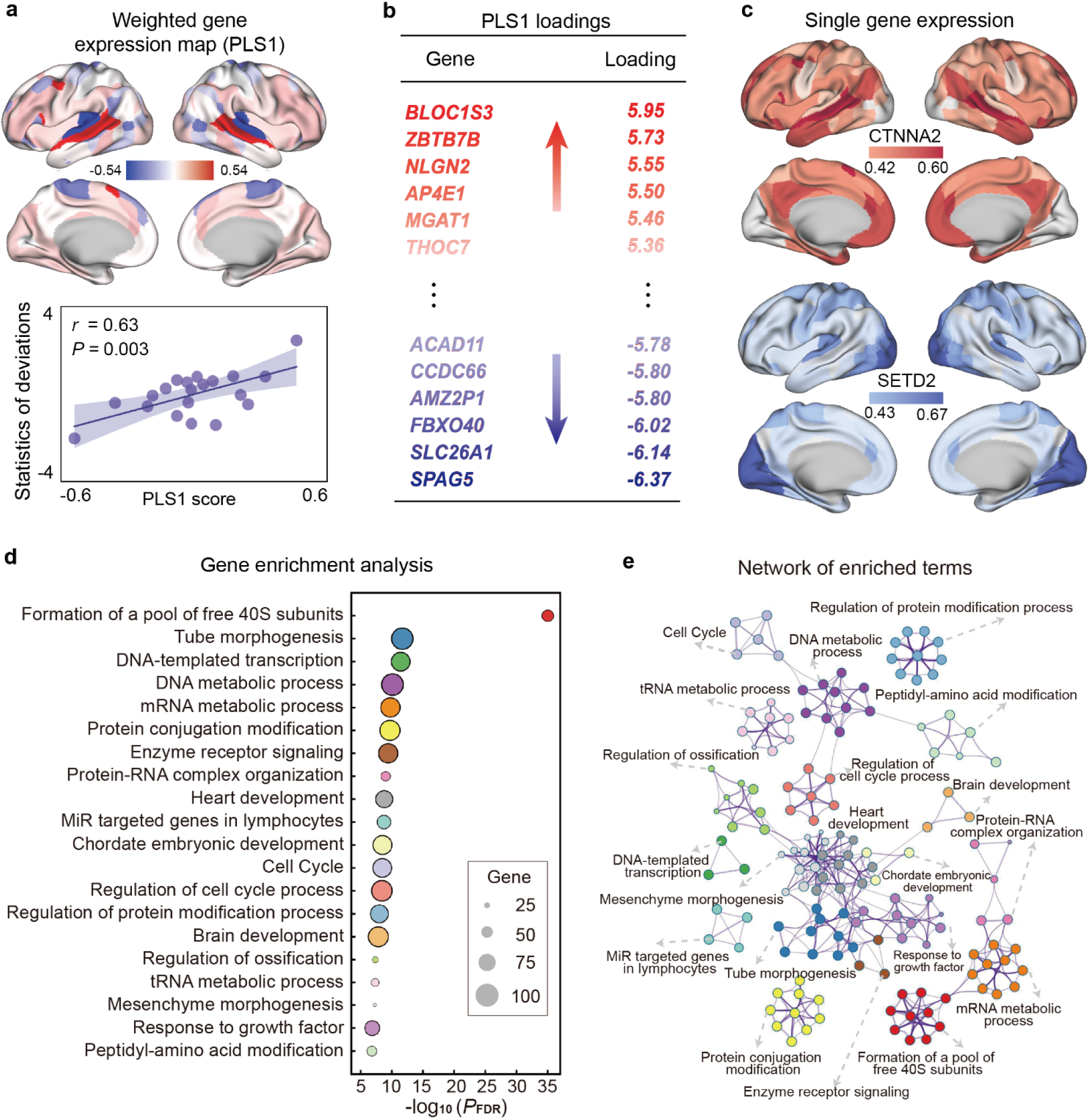
Differential gene expression profiles between participants with ASD and HCs. **a** A weighted cortical gene expression map of the regional partial least squares (PLS) scores (top panel). Scatterplot showing the relationship between weighted cortical gene expression PLS1 score and statistics of deviations between ASD and HC groups (bottom panel, *r* = 0.63, *P* = 0.003). **b** Genes were ranked on the basis of their loadings in PLS1. **c** Brain maps display the expression patterns of specific genes (*CTNNA2*, *SETD2*). Each brain map shows the expression range of a gene, with color scales indicating different expression levels. **d** Representative enriched terms of the PLS1− genes, highlighting Gene Ontology (GO) biological processes related to synaptic signaling and Reactome gene sets associated with the nervous system. The size of the circle denotes the number of genes within the given ontology term, and the multi-test with FDR corrected *P*-values are plotted as -log^10^ transformed values. **e** Metascape network plot of the enriched terms, capturing the intra-cluster and inter-cluster similarity relationships between them. Each term is represented by a circle, colored by its cluster identity and scaled proportional to the number of genes involved.

We prioritized genes based on their loadings in PLS1 (Fig. 5b), identifying top positively loaded genes (i.e., *BLOC1S3*, *ZBTB7B*, and *NLGN2*) and negatively loaded genes (i.e., *SPAG5*, *SLC26A1*, and *FBXO40*). Mutations in neuroligin genes have been linked to ASD in multiple studies^60,61^. Among the overlapping genes, *NLGN2* (Z = 5.55) as a member of the neuroligin family, has been shown in experimental mice to play a key role in maintaining the excitation–inhibition balance^62^. Furthermore, *CTNNA2* (Z = 3.33) a gene known for regulating neuronal migration and the stability of neurites^63^, has been closely associated with intellectual disability and with language development delay in patients carrying deletions involving fragment of *CTNNA2*^64,65^. This gene was highly expressed in the language network of the cerebral cortex, and its spatial distribution shows a marked overlap with the functional deviation map of ASD brains (Fig. 5c, top panel). Similarly, *SETD2* (Z = –3.04), a gene frequently mutated in ASD, may contribute to the disorder through loss-of-function effects that disrupt neuronal gene expression profiles and synaptic plasticity^66,67^, was highly expressed in frontoparietal, premotor, and visual networks (Fig. 5c, bottom panel). The top 2,000 with the highest positive Z-scores (*Z* > 1.86) were designated as the PLS+ gene set, while the 2,000 with the lowest negative Z-scores (*Z* < -2.55) were designated as the PLS− gene set. The robustness of these gene weights was supported by bootstrap-derived Z-score distributions, which showed distinct separation between the PLS+ and PLS− gene sets (Supplementary Fig. 8c). To further elucidate the biological functions of the associated genes, we performed enrichment analysis on the PLS+ and PLS− genes using Metascape^68^. Notably, the gene enrichment analysis revealed that PLS− genes were significantly associated with key terms such as “formation of a pool of free 40S subunits”, “tube morphogenesis”, “DNA-templated transcription”, and “DNA metabolic process” (Fig. 5d), suggesting their roles in maintaining genomic stability and regulating intracellular signaling. Moreover, these genes were significantly enriched in a variety of functional pathways (Fig. 5e), including Gene Ontology (GO) biological processes^69^ and Reactome pathways^70^ including “heart development”, “Protein–RNA complex organization”, and “brain development”, underscoring the integrated functions of PLS1− genes in ASD related molecular mechanisms. This gene enrichment analysis further showed that PLS+ associated genes are significantly involved in key biological processes such as “ubiquitin protein degradation”, “macroautophagy”, and also “brain development” with high statistical confidence (Supplementary Fig. 8d). Notably, enriched genes include “regulation of cell projection organization”, “metabolism of RNA”, and “regulation of autophagy” (Supplementary Fig. 8e). These findings highlight specific molecular mechanisms underlying the differences in functional network topology between individuals with ASD and HCs, suggesting that macroscale network alterations may be driven by distinct genomic regulatory pathways^71^.

## Discussion

In this study, we aimed to dissect the neurobiological heterogeneity of ASD by mapping each individual’s deviation from normative models of functional networks topology development. Using a large sample of male individuals with ASD and HCs from the ABIDE I and II datasets, we revealed striking and widespread patterns of topological deviation at individual-level in ASD. These patterns were highly heterogeneous across participants, most notably marked by an expansion of the language network, which emerged as a core system exhibiting atypical reconfiguration in individuals with ASD. Building upon these findings, we further stratified the ASD populations into three distinct subtypes, characterized respectively by perceptual and attentional dysfunctions, deficits in integrative and executive functions, and socio-emotional impairments. Moreover, by integrating transcriptomic data, we explored the genetic architecture associated with these networks with marked deviation, revealing potential genetic bases regulating the observed alterations in functional brain parcellation. These findings provide novel topological insights into the biological underpinnings of ASD and highlight the importance of linking macroscale connectomic alterations to molecular mechanisms to better understand the heterogeneity of ASD.

Precision functional mapping with ASD revealed a significant expansion of the language network. This finding aligns with previous neuroimaging research that has linked language-related cortical regions—such as the inferior frontal gyrus and superior temporal gyrus—to the neurobiology of ASD. For example, anatomical enlargement, functional abnormalities, and atypical connectivity patterns have been observed in these regions in individuals with ASD, including right superior temporal gyrus hypertrophy, atypical recruitment of extrastriate cortices during lexicosemantic processing, and disrupted early connectivity within language networks in infants at high familial risk^72–75^. Additional evidence, such as enhanced functional connectivity within language circuits^49,76^ and local cortical thickening in the left inferior frontal gyrus and superior temporal regions^77,78^, further supports the view that atypical development or delayed maturation of the language network is a core feature of ASD. Our work extends this notion by showing that network topology can act as a neural marker to reveal developmental abnormalities in ASD, with a marked expansion of language networks. This expansion appears to be primarily driven by shifts in the boundaries of the language network, often encroaching upon by the auditory network. Although more work will be required to elucidate the mechanisms underlying language network expansion in ASD, key results from previous studies indicate at least two points of view. First, converging evidence indicates that individual differences in network topology are regulated by activity-dependent mechanisms and closely linked to how frequently a given network is engaged^13^. Individuals with ASD often exhibit defects in language comprehension and expression, have difficulty organizing language that conforms to social norms, and are often accompanied by abnormal behaviors such as repetitive speech and self-talk^79–81^. In addition, it has been found that individuals with ASD are more inclined to focus on local semantic features (e.g., individual words) and less likely to integrate broader contextual information during language processing^82^. These factors may contribute to the fact that individuals with ASD involve a wider range of brain regions during language processing than is typical of the pattern in HCs. Second, during language processing, individuals with ASD may recruit a different cortical region than HCs to compensate for deficits in typical language function^83^. This alternative activation pattern may lead to an over-prioritization of language information in neural processing, which compresses the resource allocation to brain regions dependent on other tasks, ultimately leading to an atypical expansion of the brain’s language topology^84^. For example, in the ASD group, language regions showed increased connectivity with the posterior cingulate cortex and visual regions. Moreover, the posterior cingulate cortex (a core hub of the default mode network) mediated the functional connectivity between the bilateral inferior frontal and visual regions, suggesting that non-language regions are atypically integrated into the language network^45^. Thus, this functional compensatory mechanism may lead to the movement of the boundaries of the language network and thus the expansion of the language network.

Compared to the normative distributions estimated from HCs, individuals with ASD showed significant deviations in several functional networks. This finding helps explain why traditional case-control analyses often fail to detect robust differences at the group level^85,86^. By shifting toward individualized deviation modeling, we were able to capture the high degree of heterogeneity within the ASD population. Our results extended the functional connectivity based normative modeling framework by incorporating surface area percentage of individualized functional mappings into the analysis, enabling a quantitative characterization of functional brain topology. Among functional networks, the most pronounced positive deviation was the overall expansion of the language network. Meanwhile, the auditory network showed relative contraction, which may be associated with impairments in voice perception and pitch recognition commonly seen in ASD^87^. Within the somatomotor network, the somatomotor-foot region exhibited reduced surface area, which may be due to poorer motor coordination and reduced engagement in physical activities among participants with ASD, leading to decreased usage of lower limb functions and subsequent functional degradation of this area^88^. The mechanism of this topology variations also aligns with previous findings that ASD individuals show impaired performance in implicit motor sequence learning tasks, accompanied by reduced activation in the right superior parietal lobule^89^. In the visual system, the V5 area—primarily responsible for processing visual motion—also showed decreased surface area. This could be related to the visual attention biases in ASD toward static features (e.g., patterns and edges)^90,91^, with reduced focus on dynamic visual scenes, especially those involving faces and body movements in social contexts^92^. Additionally, common ASD features such as gaze avoidance and abnormal oculomotor control may further contribute to the underuse and consequent weakening of this region.

It is worth noting that although these trends of network topology are representative, we also observed both positive and negative deviations within the same networks across different individuals with ASD. This diversity further suggests that ASD is not a structurally homogeneous disorder; rather, its functional topography may be influenced by distinct neural phenotypes or underlying subtypes.

Based on the heterogeneity of deviations across ASD individuals, we established three different subtypes of ASD, which showed neurological differences in core behaviors such as social interaction, communication, and stereotypic behaviors^40,93^. Our results extend previous studies that primarily analyzed subtypes relied on clustering based on behavioral metrics or structural MRI features^94,95^. Specifically, these three subtypes were characterized by perceptual and attentional dysfunction with reduced cognitive control, deficits in integrative and executive function, and socio-emotional impairments involving affective and sensory processing, respectively^96^. Furthermore, each subtype exhibited distinct deviation patterns across large-scale brain networks, including language, default mode and sensory systems, that map onto specific behavioral phenotypes. Individuals with pronounced perceptual and sensorimotor disruptions showed difficulties in processing social cues and increased stereotyped behaviors, likely reflecting impaired visual-social integration and weakened cognitive control^97,98^. Those with deficits in integrative and executive function exhibited impairments in language use, planning, and contextual understanding, driven by disruptions in high-order control networks such as the default mode network and SCAN. In contrast, socio-emotional impairments involving affective and sensory processing were linked to reduced integration in salience and language networks, along with compensatory over-engagement of face-processing and attentional systems. These findings not only established frameworks that involved the state-of-art precision functional brain network topology, but also demonstrate the utility of combining neural deviation patterns with behavioral phenotyping to capture the multi-level heterogeneity of ASD.

Autism is widely recognized as a highly heritable condition^99,100^. Twin and family studies have demonstrated that its genetic architecture is also highly polygenic, with numerous genetic factors contributing to the disorder^32^. Despite considerable efforts to identify reliable genetic markers for autism, the diagnostic positivity rate of genetic testing is still far below the level demanded for clinical application. This limitation highlights the challenges of relying solely on genetic insights for effective clinical diagnostics^101^. In this study, we aimed to explore genetic explanations for the observed functional network deviations in ASD by taking a topological perspective and integrated these spatial maps of brain alterations with postmortem cortical gene expression profiles. This analysis identified ASD associated genes with roles in synaptic function (*NLGN2*), neuronal migration and neurite stability (*CTNNA2*), and chromatin-mediated gene regulation (*SETD2*), with *CTNNA2* showing high expression in cortical language networks and spatial overlap with ASD related functional alterations^61–67^. These findings provide a molecular perspective on functional precision mapping results and contribute to a growing body of evidence linking macroscale brain network organization to transcriptomic regulation. While prior transcriptomic studies have identified ASD-associated genes and implicated developmental pathways^102,103^, relatively few have spatially mapped gene expression onto regions exhibiting functional network topological alterations. Our study addresses this gap by applying PLS framework to spatially anchor transcriptomic patterns to functionally atypical cortical regions in ASD. The regions showing the strongest deviations also exhibited high expression of genes involved in cortical development, synaptic signaling and language processing, supporting that the functional topologic of the brain may be shaped by spatial gene expression. This approach advances previous work, rather than focusing only on case-control group differences, by linking individualized network deviations to regionally enriched molecular signals.

Despite this study used an individualized modeling framework to reveal atypical features of the functional brain atlas of ASD, several limitations need to be pointed out. First, this study is limited by the characteristics of the available datasets. The ABIDE I and II datasets were from multiple research institutions using different MRI scanners and scanning protocols, with clinical phenotyping data—including ADOS-2, SRS, and ADI-R scores—were only available for a subset of ASD participants^7^. Therefore, the associations between deviation-based subtypes and behavioral phenotypes may be biased or underestimated, as the available behavioral measures are incomplete and not uniformly collected across participants. Second, this study is based on cross-sectional characterization and lacks investigation of developmental trajectories and disease progression in longitudinal design. Compared to cross-sectional study, the longitudinal design is critical for capturing the dynamics of the distribution in brain function and structure over time, especially during critical periods of neurodevelopment^104^. However, longitudinal fMRI cohorts specifically focused on ASD remain extremely limited. To obtain a more comprehensive understanding of developmental trajectories in ASD, future research will require continuous tracking data collection, open data sharing, and multi-center collaboration for obtaining longitudinal developmental cohort in ASD. Third, our transcriptomic analysis relied on the Allen Human Brain Atlas (AHBA), which provides microarray-based gene expression profiles from 3,702 brain samples collected from only six healthy adult postmortem donors. Due to the dataset is derived from adult tissue, it cannot capture the dynamic gene regulatory processes that occur during early development—a critical period for cortical specialization and the formation of functional networks. Despite this limitation, the statistical approaches used in this study have been demonstrated to be robust in previous research linking functional connectivity differences to gene expression^105–107^. Thus, our analysis still offers a valuable foundation for future investigations, and the methods we employed have been validated in prior studies^108^.

In conclusion, our findings demonstrate that individualized precision functional mapping combined with normative models can uncover stable neurofunctional deviations, biologically grounded subtypes, and genetic correlates in ASD. This multilevel approach offers a scalable framework for understanding ASD heterogeneity, with potential implications for biomarker discovery and personalized intervention strategies.

## Methods

### Participants

In this study, we used datasets of the Autism Brain Imaging Data Exchange (ABIDE) I and II (http://fcon_1000.projects.nitrc.org/indi/abide)^109,110^. Considering the gender distribution imbalance in the incidence of ASD^111^, we included only male participants into this study. Additionally, we restricted the age range to 6–30 years, included only participants with full-scale IQ (FIQ) > 70^112^, controlled for head motion (mean framewise displacement < 0.3 mm)^113^, and ensured matching of age and head motion between the ASD and HC groups. Specifically, HCs had no psychiatric history and were age-matched to ASD individuals within each site. The final sample consisted of 554 individuals with ASD and 628 HC individuals (Table 1), recruited from 25 distinct research sites. The Autism Diagnostic Observation Schedule, Second Edition (ADOS-2)^114^ and the Social Responsiveness Scale (SRS)^115^ were used to evaluate symptom severity. Additionally, the Autism Diagnostic Interview-Revised (ADI-R), which captures developmental history and core autism symptoms through caregiver interviews was used for behavioral association analysis^116^. All studies received approval from local institutional review boards (IRBs), and the datasets were fully de-identified in compliance with HIPAA regulations, including removal of facial features from structural MRI scans^117^.

### Data acquisition and preprocessing

All resting-state fMRI (rs-fMRI) and high-resolution T1-weighted (T1w) structural MRI data were acquired using 3T MRI scanners (Siemens or Philips) across all sites (for detailed acquisition parameters, please see http://fcon_1000.projects.nitrc.org/indi/abide/)^109,110,118^. We subsequently preprocessed the data using the ABCD-HCP functional pipeline via Docker (https://hub.docker.com/r/dcanumn/abcd-hcp-pipeline)^21^. The pipeline includes motion correction, spatial distortion correction, slice timing correction, coregistration to structural images, intensity normalization, and projection of fMRI signals onto the cortical surface. We resampled rs-fMRI surface data to fs_LR32k template, a mesh from the Human Connectome Project pipeline (https://github.com/Washington-University/Pipelines). Then, prior to applying the precision functional mapping pipeline, we loaded the preprocessed CIFTI dense time series for each subject and concatenated all runs across sessions along the time axis. The concatenated series were demeaned per vertex. The preprocessed time-series data were subsequently spatially smoothed on the cortical surface using geodesic Gaussian kernels with FWHM = 2.55 mm, implemented via the Workbench tool wb_command (-cifti-smoothing, https://www.humanconnectome.org/software/workbench-command/)^12,119^. The preprocessed time series data is used for subsequent individualized brain functional mapping.

### Individualized functional brain parcellation

For each participant, the denoised rs-fMRI time series data which mapped to the individual’s fs_LR32k cortical surface were used to compute a distance matrix based on the left and right hemisphere on fs_LR32k midthickness surfaces. Then, an individual-level functional connectivity matrix was constructed by computing the Pearson’s correlations between the time courses of all cortical vertices. To minimize the influence of spatial proximity and enhance reliabilities, the top 0.1% of the strongest connections for each participant were retained to focus on the most reliable interactions^13^.

The thresholded matrices were then put into the Infomap algorithm with fixed free parameters (for example, the number of algorithm repetitions) across all subjects to ensure consistency. The Infomap algorithm, which is a community detection method based on information theory, was used for functional community clustering^120^. The optimal scale for each individual was defined as the threshold that produced the best size-weighted average homogeneity, which maximized the median of the size-weighted average uniformity calculated with respect to random network rotations^121^. The Infomap algorithm identified a variable number of functional communities per subject, typically ranging from approximately 60 to 200, using fixed free parameters. Each resulting community was assigned to one of 20 canonical functional networks (default-parietal, default-anterolateral, default-dorsolateral, default-retrosplenial, frontoparietal, dorsal attention, language, salience, cingulo-opercular, parietal memory, SCAN, visual V5, visual stream, lateral visual, visual V1, premotor, somatomotor-hand, somatomotor-face, somatomotor-foot, auditory) based on its spatial location and functional connectivity pattern with the same predefined reference atlas^13^.

Then, we used the wb_command (-surface-vertex-areas) to calculate the surface area (in mm^2^) of each vertex on the midthickness cortical surface for each subject. Next, by dividing the total surface area of all vertices within a specific functional network by the total surface area of the entire cortex, we obtained the proportion of the network in the cortex, which was used as an indicator to measure the distribution range of the network topology in the subsequent analysis. Considering that the data were collected from 25 different sites, we applied ComBat harmonization to mitigate non-biological variability introduced by site-specific factors, while preserving meaningful biological differences^41,42^. The effectiveness of the harmonization was assessed using analysis of variance (ANOVA).

### Quantification of inter-individual spatial variability

To assess the spatial variability of functional network topography across individuals, we computed the Dice coefficient for each pair of subjects within the ASD and HC groups across 20 canonical networks. The Dice coefficient was calculated by quantifying the spatial overlap of each network between pairs of individual functional atlases:

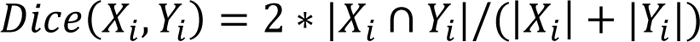

where *X*_*i*_and *Y*_*i*_ denote the set of surface vertices assigned to network *i* in participants *X* and *Y*, respectively, and ∩ indicates the intersection of these vertex sets within each group (ASD and HC). The Dice coefficient ranges from 0 (no overlap) to 1 (perfect overlap), reflecting the spatial consistency of a given network between two individuals^37^. The distributions of Dice coefficient for each network were then statistically compared using a two-sample Kolmogorov–Smirnov (KS) test between ASD and HC groups.

To further quantify inter-individual heterogeneity in cortical organization, we calculated the coefficient of variation (CV = standard deviation / mean) of Dice coefficient for each network, and then applied a general linear model (GLM) to regress out the group-average network size within the ASD and HC groups. The CV accounts for baseline differences in average network size and provides a normalized measure of inter-individual variability in functional network organization^46^.

### Constructing normative models

To construct the normative trajectories of various functional precision mapping metrics in healthy individuals based on surface area percentage, we applied the generalized additive models (GAMs). The GAMs extend generalized linear models by allowing non-linear functions of the predictors while maintaining additive structure and interpretability^122^. Therefore, to avoid the normative model’s being biased by individuals with ASD, we used GAMs to estimate the normative surface area percentage for each of the brain networks defined by functional precision mapping only based on the data from HCs. Specifically, for each region *i*, the surface area percentage was modeled as a smooth function of age *f*_1_(*Age*), with mean head motion included as a linear covariate. The model takes the following form:

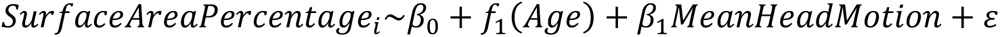

where *β*_0_ is the intercept, *f*_1_(*Age*) represents the nonliear age effect estimated by GAMs, *β*_1_ is the coefficient of mean head motion, and *ɛɛ* is the residual error term. To ensure the generalizability and robustness of the model, we performed 10-fold cross-validation within the model.

To estimate individual deviations, the *i*th regional surface area percentage of each individual with ASD were mapped onto the normative models derived from HCs corresponding to their age. We defined a z-value for each brain region to quantify deviation from the normative model, based on the difference between the observed surface area percentages and the model-predicted values:

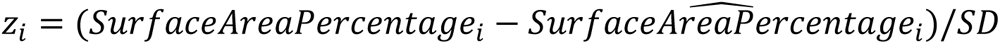

Specifically, to identify extreme deviations in ASD patients, we set a threshold of |*z*| > 2.6, corresponding to the 5th and 95th percentiles of the normative distribution.

### Subtypes of ASD and associations with behaviors

Based on the normalized model, we estimated the deviation in surface area percentage for 20 canonical brain networks in each individual with ASD. Then, we employed a data-driven k-means clustering algorithm to explore ASD subtypes characterized by distinct deviation patterns, with the optimal number of clusters (k = 3) determined at the inflection point of the elbow method curve^40^ and successfully validated by multiple classifiers. To identify latent relationships between ASD subtypes and behavioral measures, we used Canonical Correlation Analysis (CCA), a multivariate statistical method that identifies linear combinations of variables in networks surface area and behavioral scales with maximal correlation^123^. To address the issue of multicollinearity, we incorporated an L2 regularization term (*r* = 0.1) into the CCA framework, resulting in the regularized CCA (rCCA) ^124,125^:

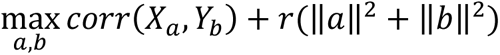

where *X* and *Y* represent the functional network topological and behavioral feature matrices, respectively, while *a* and *b* are their corresponding projection vectors.

We utilized three widely recognized ASD-related instruments to quantify behavior, including the ADOS-2, SRS, and ADI-R. From these instruments, we selected subscales that specifically focus on social and communication impairments and stereotyped behaviors. We then conducted 1000 permutation tests within each ASD subtype, randomly shuffling behavioral data while keeping functional network topological data fixed to construct a null distribution of canonical correlations and calculate empirical *P*-values for assessing statistical significance.

### Classification analysis

To assess whether individualized deviations across functional brain networks could reliably predict ASD subtypes, we implemented a series of supervised classification framework using Python’s scikit-learn library (version 1.2.2). A total of 20 canonical network’s deviation features were extracted for each participant, corresponding to the z-normalized deviation values within each of the 20 canonical functional networks. Prior to classification, missing values were imputed using the mean across individuals, and all features were standardized using z-score normalization^126^. We evaluated the classification performance of three widely used classifiers, including random forest (RF), support vector machine (SVM) with a linear kernel, and logistic regression (LR) with L2 regularization.

To preserve subtype proportions, the dataset was stratified into a training set (70%) and a testing set (30%), and this stratified splitting procedure was randomly repeated 10 times. Within each training set, 5-fold cross-validation was performed to optimize hyperparameters for each model. Classification performance was assessed using accuracy on the test set, and model predictions were further evaluated using confusion matrices and classification reports, including precision, recall, and F1-score metrics. To handle the multiclass nature of ASD subtype prediction, we adopted a one-against-one strategy for the SVM classifier, where binary classifiers were trained between each pair of subtypes and the final prediction was determined by majority voting, with linear kernel was used, and the regularization parameter *C* set to the default value of 1.0. For the RF classifier, we tuned the number of decision trees and their maximum depth, and the final model included 100 trees with the default depth setting. Feature importance scores were extracted to identify the most discriminative brain networks. LR was implemented with L2 regularization, and the maximum number of iterations was increased to 1000 to ensure convergence.

### Genetic association analysis

We employed cortical gene expression data from the Allen Human Brain Atlas (AHBA, http://human.brain-map.org), which contains postmortem microarray measurements from six adult donors (age range: 24–57 years, one female) across 3,702 sampling sites^127,128^. The preprocessing of gene expression data followed the procedure outlined by Markello et al.^129^, and consisted of the following steps: (i) updating the probe-to-gene annotations, applying intensity-based filtering with a threshold of 0.5, and selecting the probes that exhibited the most consistent regional expression patterns across donors; (ii) mapping the samples to a group-level 20-region parcellation template derived from the individual-mode consensus using a bilateral mirroring approach, and applying centroid-based imputation for regions with missing data; (iii) normalizing the gene expression values within each donor, first across genes and then across samples; (iv) averaging the expression values of corresponding regions across all donors to generate a 20 × 15,632 gene expression matrix for subsequent analyses.

To investigate the transcriptomic correlates of ASD-related deviations in brain functional organization, we applied partial least squares (PLS) regression to relate the Z-normalized gene expression data with the regional surface area percentage deviations between individuals with ASD and HCs across 20 canonical brain networks^59^. Specifically, the first PLS component (PLS1) was extracted to capture the dominant gene expression profile associated with the spatial distribution of individualized brain deviations in ASD. The significance of the explained variance by PLS1 was assessed using 1,000 spatial spin permutations to correct for spatial autocorrelation^130^. Bootstrapping was conducted to estimate standard errors of gene weights, and Z-scores were computed for ranking genes based on their contributions to PLS1.

Genes were ranked based on their normalized PLS1 weights, and the top 2,000 with the highest positive Z-scores (Z > 1.86) were designated as the PLS+ gene set, while the 2,000 with the lowest negative Z-scores (Z < -2.58) were designated as the PLS− gene set. Each gene set was submitted to Metascape (https://metascape.org), an automated tool for enrichment analysis of biological pathways^68^. Significance of enriched terms was determined with a false discovery rate (FDR) threshold of *q* < 0.05.

## Data availability

The raw and pre-processed data are available from the Autism Brain Imaging Data Exchange (ABIDE) I and II (http://fcon_1000.projects.nitrc.org/indi/abide/) and Allen Human Brain Atlas (AHBA, http://human.brain-map.org). Numerical source data generated in this study are included in Supplementary Data.

## Author contributions

G.Y. and R.Y. designed research; R.Y. analyzed the data; R.Y. and G.Y. wrote the first draft of the manuscript; X.W., S.L., J.L., Z.W., W.Z., T.G. and G.Y. edited the manuscript.

## Competing interests

The authors declare no competing interests.

## Notes

### Competing Interest Statement

The authors have declared no competing interest.

